# Atypical retinal function in a mouse model of Fragile X syndrome

**DOI:** 10.1101/2024.03.15.585283

**Authors:** Anna L. Vlasits, Maria Syeda, Annelise Wickman, Pedro Guzman, Tiffany M. Schmidt

## Abstract

Altered function of peripheral sensory neurons is an emerging mechanism for symptoms of autism spectrum disorders. Visual sensitivities are common in autism, but whether differences in the retina might underlie these sensitivities is not well-understood. We explored retinal function in the Fmr1 knockout model of Fragile X syndrome, focusing on a specific type of retinal neuron, the “sustained On alpha” retinal ganglion cell. We found that these cells exhibit changes in dendritic structure and dampened responses to light in the Fmr1 knockout. We show that decreased light sensitivity is due to increased inhibitory input and reduced E-I balance. The change in E-I balance supports maintenance of circuit excitability similar to what has been observed in cortex. These results show that loss of Fmr1 in the mouse retina affects sensory function of one retinal neuron type. Our findings suggest that the retina may be relevant for understanding visual function in Fragile X syndrome.

## INTRODUCTION

Global symptoms of autism spectrum disorders (ASD) likely arise from changes in neural function in many different modules of the brain. Recent findings suggest that some symptoms of ASD stem from atypical sensory processing (Robertson and Baron-Cohen 2017; Falck-Ytter and Bussu 2023), including hyperexcitable peripheral nerves (Orefice et al. 2016; McCullagh et al. 2020). However, research into vision in ASD has primarily focused on the cerebral cortex (Simmons et al. 2009; Robertson and Baron-Cohen 2017), with less evaluation of earlier processing stages, such as in the retina. ASD has many symptoms that could be related to altered function of the retina. For example, in Fragile X syndrome, a disorder strongly linked to ASD, symptoms include reduced temporal resolution of vision in infants (Farzin, Rivera, and Whitney 2011), lower contrast sensitivity (Kogan et al. 2004), visual hypersensitivity (Raspa et al. 2018), and sleep disturbances such as night waking (Hagerman et al. 2017). Some of these symptoms are homologous in the mouse model of Fragile X syndrome, which supports the idea that “low level” retinal processing may shape symptoms (Saré et al. 2017; Goel et al. 2018; Perche et al. 2021; Felgerolle et al. 2019; Yang et al. 2022).

The retina is a multi-layered set of neural circuits with intricate wiring of over 100 different cell types (Vlasits, Euler, and Franke 2019; Baden et al. 2018), supporting not only our conscious visual experience, but also reflexive and non-reflexive eye movements, our circadian rhythms, and our mood and affect (LeGates, Fernandez, and Hattar 2014). These different roles are accomplished by neural circuits in the retina that sort different channels of light information—such as motion, contrast, and time of day—and then relay that information onward. These final output channels of the retina are different types of retinal ganglion cells, which are axon-bearing neurons that project from the eye to many different brain areas (Kerschensteiner 2022).

Loss of ASD-linked genes across the brain leads to changes in an array of cellular properties, including anatomical and physiological changes in excitatory and inhibitory neurons (Contractor, Ethell, and Portera-Cailliau 2021; Zhao et al. 2022). Intriguingly, many ASD-linked genes are also expressed in the retina (**Fig. 1A**), where excitatory and inhibitory interneurons shape ganglion cells’ tuning for specific types of light information. Gross physiology of the human retina in people with ASD using electroretinography has revealed slower and lower amplitude responses of the optic nerve, suggesting that atypical visual processing in ASD begins in the retina (Perche et al. 2021; Constable et al. 2020). However, whether and how the function of individual types of retinal ganglion cells is altered in ASD has not, to our knowledge, been evaluated.

**Figure 1.**
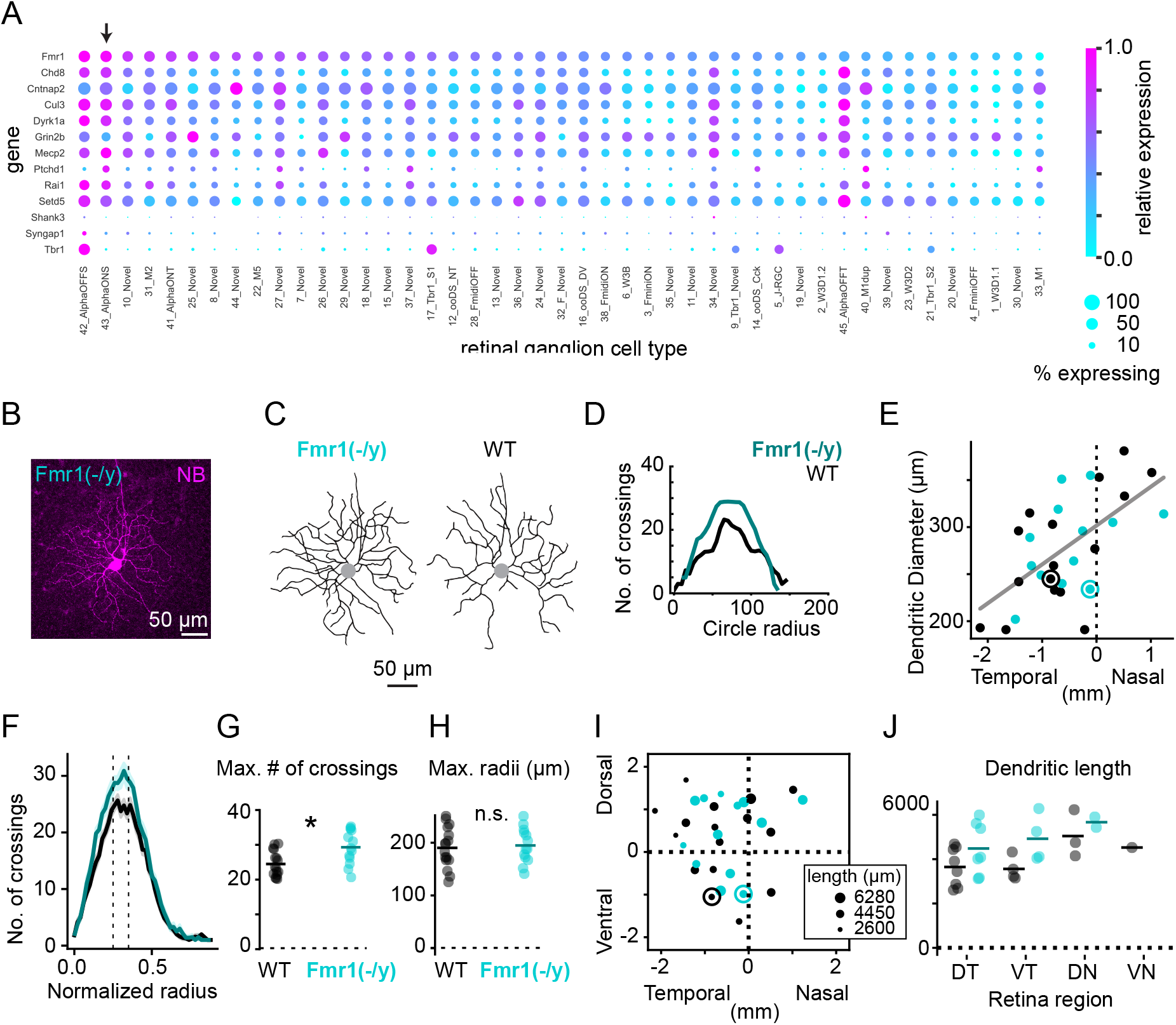
sOn-α retinal ganglion cells have denser dendrites in Fmr1(-/y) mice. A) Type-specific expression of ASD-linked genes in mouse retinal ganglion cells from a published transcriptome dataset (Tran et al. 2019). Color: relative expression level within each gene. Dot size: percent of cells of each type expressing that gene. Arrow: the sOn-α RGC. B) Z-projection of a confocal image showing neurobiotin-filled dendrites of an sOn-α cell in an Fmr1(-/y) retina. C) Traced skeletons of two example cells from an Fmr1(-/y) mouse and WT mouse. D) Sholl analysis of the example cells in **C**. E) Dendritic diameter of a population of sOn-α cells plotted as a function of the nasal-temporal position on the retina. Dots with circles around them: example cells from **C**. Grey line: linear regression to the entire population, R-squared = 0.37. F) Mean sholl analysis for populations of sOn-α cells from WT and Fmr1(-/y) retinas. Shaded regions: standard error. Dotted lines: region used for analysis in **G**. G) Max. number of crossings in sOn-α cells from WT and Fmr1(-/y) mice. Dots are individual cells, line is the mean. * indicates statistical significance. H) Maximum radii of sOn-α cells from WT and Fmr1(-/y) mice. N.s. = not significant. I) sOn-α cells plotted on their retina coordinates, with dot size indicating the total dendritic length. Circled dots: example cells from **C**. J) Total dendritic lengths of cells within each retinal quadrant: dorsotemporal (DT), ventrotemporal (VT), dorsonasal (DN), and ventronasal (VN).

Here, we examined ASD-linked gene expression in the mouse retina and found widespread and type-specific expression of ASD-linked genes in retinal ganglion cells, leading us to hypothesize that retinal ganglion cell function may be altered in mouse models where ASD-linked genes are disrupted. To test this, we assessed retinal ganglion cell morphology and function in a mouse knockout of an ASD-linked gene, the Fmr1 knockout model of Fragile X syndrome. We chose to measure the function of the “sustained On alpha” retinal ganglion cell (sOn-α cell) because we observed that it expresses many ASD-linked genes, including Fmr1. Using a combination of electrophysiological recordings and computational modeling, we find that changes in cellular morphology as well as a shift in E-I balance alters the signaling in sOn-α cells while maintaining stable post-synaptic potentials. These findings indicate that some visual changes in Fragile X syndrome and ASD may arise in the retina, at the earliest stages of visual processing.

## RESULTS

### sOn-*α* cells have denser dendrites in Fmr1(-/y) mice

To examine which neuronal cell types in the retina might be affected by loss of ASD-linked genes, we examined previously published transcriptomic data from the mouse retina (Shekhar et al. 2016; Tran et al. 2019; Yan et al. 2020; Goetz et al. 2022). We identified a list of ASD-linked genes from the literature and explored the mRNA expression of those genes across retinal cell types (**Figure 1A, Fig. S1**). We noted from this analysis that one ASD-linked gene, Tbr1, has already been identified as a regulator of type-specific retinal ganglion cell (RGC) development (Liu et al. 2018). We also noted widespread expression of Fmr1 across cell classes and in both mice and primates. Fmr1 is the gene involved in Fragile X syndrome, the most common genetic cause of autism. Overall, we observed type-specificity in the relative amount of expression of each ASD-linked gene and percent of cells expressing a given gene, with retinal ganglion cells and amacrine cells expressing more autism linked genes than bipolar cells. This suggests that loss of ASD-linked genes across retinal cell types may have differential effects on the function of the retina.

Based on this survey, we selected a single ganglion cell type to study further in the context of the loss of a single ASD-linked gene. We chose the sustained-On α retinal ganglion cell (sOn-α cell; arrow in **Fig. 1A**) (Krieger et al. 2017) because this type exhibited relatively high expression of most ASD-linked genes, including Fmr1 (**Fig. 1A**). In addition, the sOn-α cell is a well-studied RGC type with identified roles in visual behaviors (Kim et al. 2020; Johnson et al. 2021; Schmidt et al. 2014), which is easy to identify due to its large cell soma (Bleckert et al. 2014). We studied the sOn-α cell in the Fmr1 knockout model of Fragile X syndrome because previous work has demonstrated reduced b-wave of the electroretinogram in both mice and humans with Fragile X syndrome, which indicates altered signaling in the inner retina (Rossignol et al. 2014; Perche et al. 2021).

To determine whether loss of Fmr1 affects development of sOn-α cells, we analyzed the dendritic morphology of sOn-α cells in isolated retinas from wildtype (WT) and Fmr1(-/y) mice. We identified sOn-α cells based on physiological and molecular features that are unique to this cell type (see Methods and **Fig. S2**) (Contreras et al. 2023) and filled these cells with neurobiotin to visualize their dendrites (**Fig. 1B-C**). sOn-α cells in Fmr1(-/y) retinas show denser arbors as measured by Sholl analysis, even though their dendritic diameters were not different between genotypes (**Fig. 1D-F**, n = 16 WT cells from 10 mice, 13 Fmr1(-/y) cells from 5 mice). The maximum number of crossings was higher (**Fig. 1G**; WT: 24 ± 3 crossings, Fmr1(-/y): 29 ± 5 crossings, p = 0.005) and the total dendritic length was higher in Fmr1(-/y) retinas compared to WT (**Fig. 1I-J**; WT: 3942 ± 923 µm, Fmr1(-/y): 4799 ± 1043 µm; p = 0.032), while the maximum radii were not significantly different between the two populations of cells (**Fig. 1H**; WT: 182 ± 36 µm, Fmr1(-/y): 186 ± 30 µm). Overall, these results suggest that sOn-α cell in the Fmr1(-/y) mouse are developing denser dendrites within overall typical arbor sizes, which could lead to altered synaptic wiring and function of these cells.

### Reduced spiking responses to dim light flashes in sOn-*α* cells

The increased dendritic density of sOn-α cells in Fmr1(-/y) mice prompted the hypothesis that sOn-α cells might have altered responses to light. To test this, we performed cell-attached recordings of sOn-α cells and presented dim-photopic green-spectrum LED light (wavelength 560 nm, “dim green light”) to measure rod and cone-mediated light responses (**Fig. 2A**). Because sOn-α cells vary in their dendritic field size and visual response properties depending on their location in the retina (Sonoda, Okabe, and Schmidt 2020; Bleckert et al. 2014; Szatko et al. 2020) (see **Fig. 1E**), we confined our recordings to the temporal half of the retina. As suggested by their name, sOn-α cells in WT retinas exhibit an initial peak response to an increase in luminance (“On” response) followed by sustained firing for the duration of the light stimulus (**Fig. 2B-C**, n = 13 WT cells from 4 mice, 10 Fmr1(-/y) cells from 3 mice). Fmr1(-/y) cells still exhibited an overall On-sustained firing pattern and had similar baseline firing rates to WT (WT: 28 ± 21 Hz, Fmr1(-/y): 38 ± 21 Hz; two-sided t-test: p = 0.276). However, during the light response, the Fmr1(-/y) cells had lower firing rates compared to WT cells during both the peak (WT: 162 ± 78 Hz, Fmr1(-/y): 99 ± 62 Hz; two-sided t-test: p = 0.038) and sustained (WT: 62 ± 29 Hz, Fmr1(-/y): 29 ± 23 Hz; p = 0.005) period of the response (**Fig. 2D**). Lower firing rates were observed in the Fmr1(-/y) cells compared to WT in both the dorsal and ventral retina (**Fig. 2E-F**), despite differences in green light sensitivity between these retinal regions (Szatko et al. 2020). These results demonstrate that the sOn-α cell light response is weaker in Fmr1(-/y) mice, and points to decreased excitation and/or increased inhibition in sOn-α cells of Fmr1(-/y).

**Figure 2.**
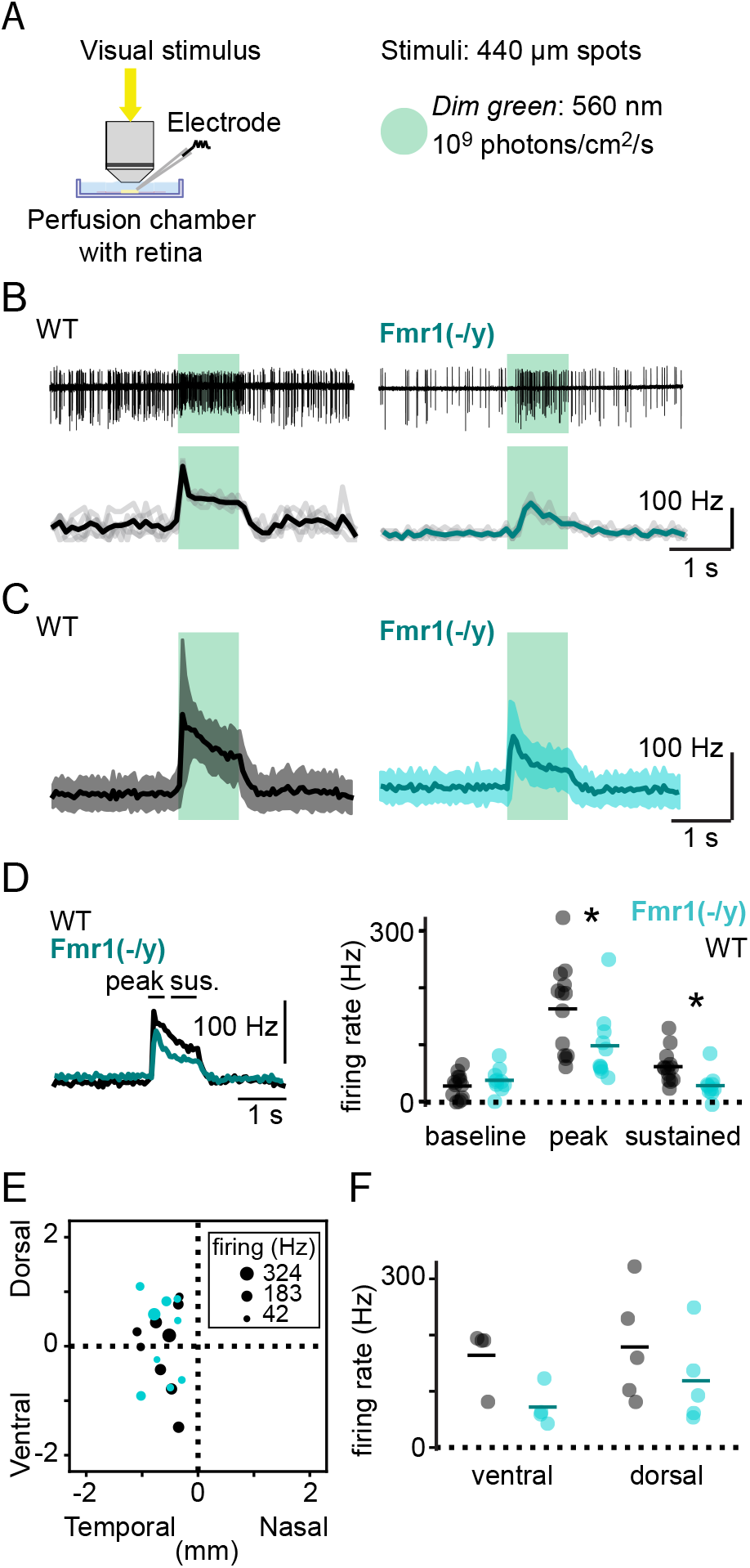
sOn-α cells have reduced responses to dim light flashes. A) Experimental setup showing microscope objective, electrophysiology electrode, perfusion chamber with the retina, and the path of the visual stimulus. Dim green stimulus was an LED source with the listed wavelength and intensity. B) Cell-attached responses of sOn-α cells to dim green light. Left: wild type (WT), right: Fmr1(-/y) knockout. Top: example recordings. Bottom: firing rate of four repetitions (light grey) and their mean (black). Timing of stimulus: green boxes. C) Population mean of firing rates for WT and Fmr1(-/y) cells. Shaded region is s.d. D) Left: population mean firing rates for WT (black) and Fmr1(-/y) (turquoise) overlaid to indicate regions used to calculate summary data. Right: baseline, peak, and sustained firing rates for individual cells (dots) and the mean (lines). * indicates significantly different means. E) sOn-α cells plotted on their retinal coordinates, with dot size indicating the peak firing rate. All cells were located in the temporal retina. F) Peak firing rate as a function of retinal location.

### Increased synaptic inhibition to sOn-*α* cells in Fmr1(-/y) mice

The reduced light sensitivity of sOn-α cells in Fmr1(-/y) mice could be the result of changes in a variety of different physiological properties of these cells. We examined whether differences in excitatory or inhibitory inputs could contribute to reduced firing in response to light flashes. We performed voltage clamp recordings from sOn-α cells to extract the time course of the excitatory (Ge) and inhibitory (Gi) conductances (**Fig. 3A-B**). We observed that while Ge was largely similar in WT and Fmr1(-/y) cells, Gi in the Fmr1(-/y) cells was significantly larger than the typical inhibitory conductance in WT (**Fig. 3B-D**; n = 8 WT cells from 5 mice, 11 KO cells from 7 mice; for Ge, WT: 171 ± 97 nS*s, Fmr1(-/y): 277 ± 152 nS*s; p = 0.083; for Gi, WT: 282 ± 264 nS*s, Fmr1(-/y): 920 ± 381 nS*s; p = 0.0004). The difference in Gi persisted throughout the time course of the response, with both the transient (**Fig. 3E**; WT: 5.30 ± 4.21 nS, Fmr1(-/y): 15.46 ± 6.15 nS; p = 0.0005) and sustained (WT: 2.53 ± 2.44 nS, Fmr1(-/y): 8.69 ± 4.03 nS; p = 0.0007) periods exhibiting a significantly greater inhibitory conductance in Fmr1(-/y) cells compared to WT. We also observed a small, but still significant, increase in the excitatory conductance in sOn-α cells in Fmr1(-/y) retinas, which was restricted to the sustained (WT: 1.36 ± 0.85 nS, Fmr1(-/y): 2.51 ± 1.41 nS; p = 0.041) and off (WT: -1.10 ± 0.77 nS, Fmr1(-/y): 0.39 ± 1.08 nS; p = 0.003) portions of the response. Most sOn-α cells in the Fmr1(-/y) exhibited larger inhibitory conductances compared to excitatory conductances in response to light flashes (**Fig. 3F**), suggesting that Fmr1(-/y) sOn-α cells may have altered excitatory-inhibitory (E-I) ratio. In addition to examining synaptic input, we measured intrinsic properties including excitability, spike properties, and the intrinsic photoresponse, and found that they were largely similar between WT and Fmr1(-/y) sOn-α cells, with a small difference in the intrinsic photoresponse (**Fig. S3**). Overall, these results suggest that the decreased spiking in response to light flashes in sOn-α cells in the Fmr1(-/y) mice is due to increased inhibitory input to these cells. Altered E-I balance is a physiological change commonly described in models of ASD (Sohal and Rubenstein 2019; Rubenstein and Merzenich 2003). To explore this further, we calculated the E-I ratio of the light response (**Fig. 3G**; see Methods). We found that the E-I ratio was lower (indicating more inhibition relative to excitation) during both transient (WT: 0.57 ± 0.03, Fmr1(-/y): 0.51 ± 0.08; p = 0.034) and sustained (WT: 0.47 ± 0.05, Fmr1(-/y): 0.36 ± 0.09; p = 0.007) periods of the response to the light flash (**Fig. 3H**). The decreased E-I ratio in sOn-a cells suggests that there may be an increased number of inhibitory synaptic inputs onto these cells in Fmr1(-/y) retinas and consequent increase in the frequency of spontaneous currents. Spontaneous synaptic activity is very high in retinal neurons, so we could not isolate individual miniature currents in our recordings. As a measure of spontaneous activity, we therefore quantified spontaneous activity by measuring the standard deviation of the spontaneous currents (Vlasits et al. 2014) (n = 11 WT cells from 5 mice, 12 KO cells from 7 mice). We found that spontaneous excitatory currents were not significantly different in WT vs. Fmr1(-/y) mice (WT: 50.91 ± 15.05 pA, Fmr1(-/y): 44.82 ± 23.18; p = 0.464), but that spontaneous inhibitory currents had higher standard deviations in Fmr1(-/y) cells compared to WT cells (**Fig. 3I-J**; WT: 35.44 ± 14.86 pA, Fmr1(-/y): 71.72 ± 41.35; p = 0.013). These results suggest that there may be more inhibitory synapses on sOn-α cells in Fmr1(-/y). This interpretation is supported by a decrease in input resistance in sOn-α cells (**Fig. 3K**; n = 18 WT cells from 8 mice, 23 KO cells from 9 mice; Rin, WT: 164 ± 63 MΩ, Fmr1(-/y): 113 ± 46; p = 0.007; Cm, WT: 26 ± 16 pF, Fmr1(-/y): 22 ± 11; p = 0.464), which would be expected in the presence of increased inhibitory input and opening of chloride channels. Overall, our results suggest that loss of Fmr1 in the retina alters sOn-α cells, leading to changes in their excitatory and inhibitory synaptic inputs.

**Figure 3.**
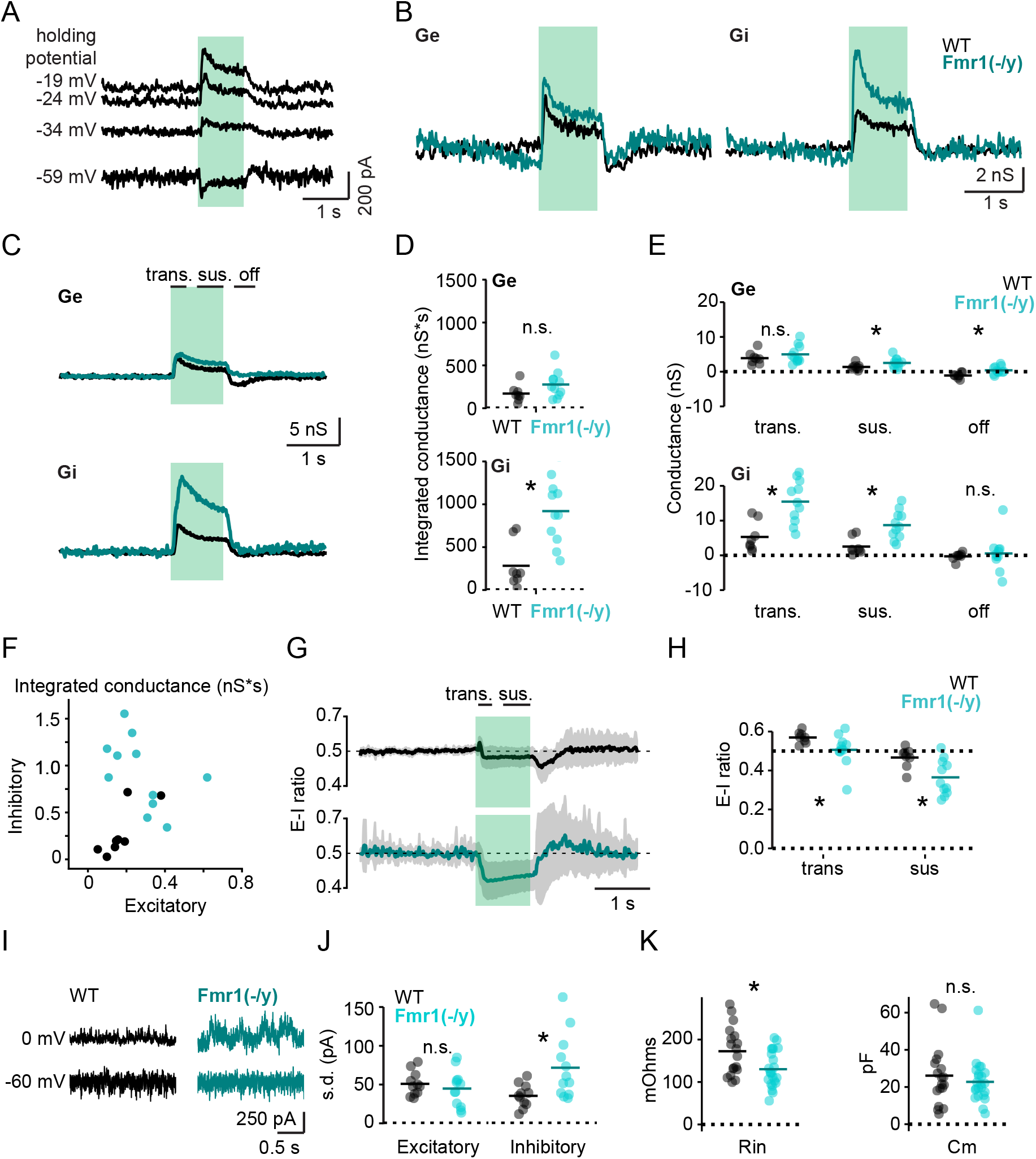
Heightened synaptic inhibition in sOn-α cells from Fmr1(-/y) mice. A) Voltage clamp recordings from an example sOn-α cell in an Fmr1(-/y) retina showing response to dim green light stimulus (green rectangle) at multiple holding potentials. B) Results of conductance analysis for two example sOn-α cells from WT (black) and Fmr1(-/y) (teal) retinas. Left: excitatory conductance (Ge), right: inhibitory conductance (Gi). C) Mean Ge and Gi for a population of sOn-α cells from WT and Fmr1(-/y) retinas. Epochs for transient (“trans.”), sustained (“sus.”), and light off (“off”) periods are indicated. Lines: group means. D) Integrated Ge (top) and Gi (bottom) for WT and Fmr1(-/y) sOn-α cells. N.s. = not significant, * = statistically significant. E) Summary values for Ge (top) and Gi (bottom) in WT and Fmr1(-/y) cells. “Trans.”: max. average conductance during the transient period. “Sus.”: mean during the sustained period. “Off”: mean during the off period. F) Relationship between integrated Ge and integrated Gi in WT and Fmr1(-/y) cells. G) The mean excitatory-inhibitory ratio (E-I ratio) over the time course of the stimulus for WT and Fmr1(-/y) cells. H) Population summary of E-I ratio during transient and sustained periods of the stimulus response. I) Spontaneous activity during the baseline period in example sOn-α cells from WT and Fmr1(-/y) retinas at holding potentials that isolate excitatory (-60 mV) and inhibitory (0 mV) currents. J) Standard deviation (s.d.) of spontaneous excitatory and inhibitory recordings in sOn-α cells from WT and Fmr1(-/y) retinas. K) Input resistance (Rin) and membrane capacitance for sOn-α cells in WT and Fmr1(-/y) retinas.

### Conductance model predicts reduced E-I ratio to stabilize post-synaptic potentials

We found that sOn-α cells in the Fmr1(-/y) mouse have larger inhibitory conductances and lower E-I ratio than WTs. This result contrasts with changes observed in other areas of the brain, where E-I ratio is often found to be higher in ASD models (Sohal and Rubenstein 2019; Contractor, Klyachko, and Portera-Cailliau 2015). Recently, Antoine et al. (2019) proposed that an increased E-I ratio may serve as a compensatory mechanism to stabilize spiking in the cortex. Using a parallel conductance model, they demonstrated that, as excitatory and inhibitory conductances scale down, E-I ratio must increase to maintain stable post-synaptic potentials (PSP).

We explored whether this model fits with our data by replicating the parallel conductance model using the average excitatory and inhibitory conductances from WT sOn-α cells (**Fig. 4A-B**). First, we compared the “Native” WT conductance scaling to two alternate cases: “Stable” scaling, in which E-I ratio is maintained with both Gex and Gin scaled up 3x; and “Decreased” scaling, in which E-I ratio decreases through scaling Gin by 3x and Gex by 1.5x, as more typically observed in our data for Fmr1(-/y) cells (**Fig. 4C**). Our model predicts that the PSPs in the “Native and “Decreased” models have similar peak amplitudes, while the “Stable” scaled model has a higher peak amplitude (**Fig. 4D**). We explored the parameter space of conductance scaling and found that when both Gex and Gin are scaled up, E-I ratio must decrease to maintain a stable PSP (**Fig. 4E**). Thus, the parallel conductance model provides evidence of a consistent trend in E-I ratios observed in cortex and in the retina in ASD models, regardless of whether excitation and inhibition are scaling up or down.

**Figure 4.**
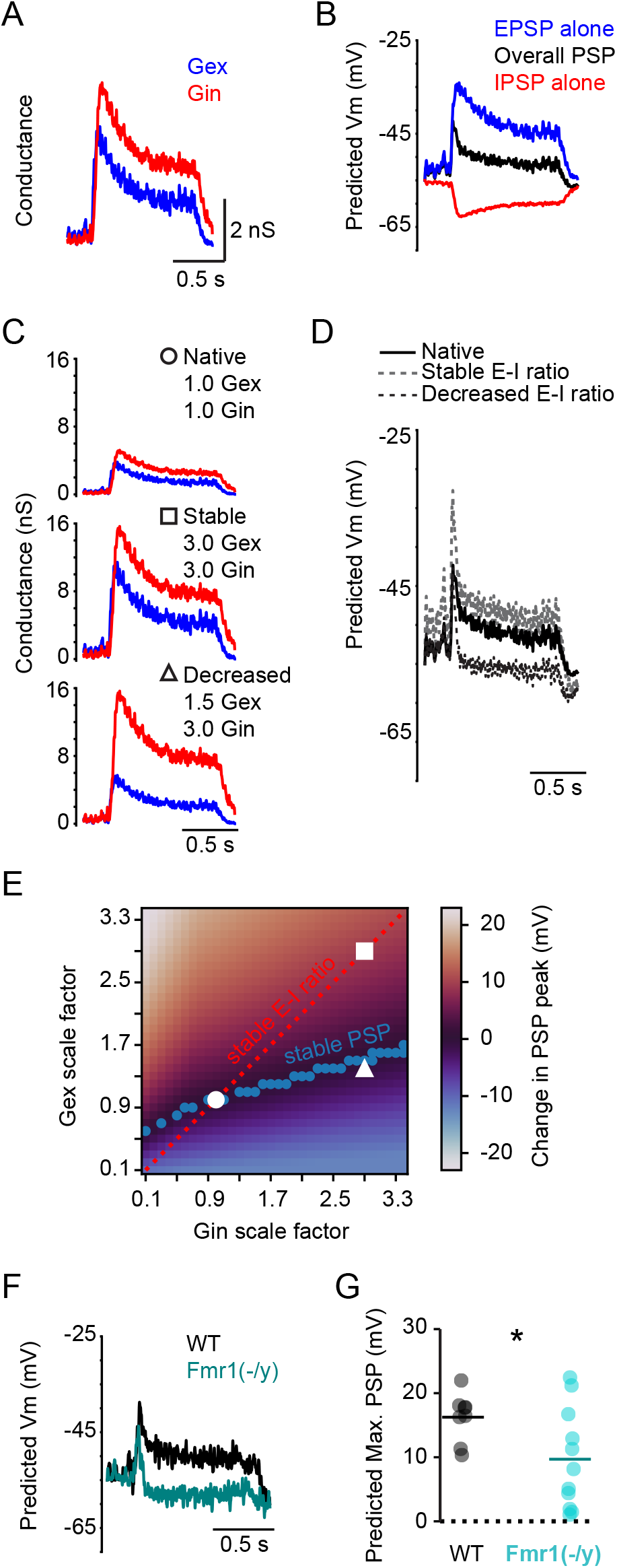
Conductance model predicts reduced E-I ratio to stabilize post-synaptic potentials. A) Average excitatory (Gex) and inhibitory (Gin) from WT sOn-α cells used in conductance model. B) Predicted membrane voltage from conductance model for three conditions: the excitatory post-synaptic potential (EPSP) predicted from Gex, the inhibitory post-synaptic potential (IPSP) predicted from Gin, and the overall post-synaptic potential (PSP) using both Gex and Gin. C) Native and scaled excitatory and inhibitory conductances. “Stable” E-I ratio scales the conductances by the same scaling factor (3x), while “decreased” E-I ratio scales up Gin (3x) compared to Gex (1.5x). D) Model prediction for the membrane voltage Vm for each of the conductance conditions in **C**. E) Heat map of the change in the post-synaptic potential (PSP) peak for different scaling factors of Gex and Gin compared to the Native case. The three conditions in **C** are shown with white symbols. Red dotted line: stable E-I ratio. Blue dots: where PSP is less than 0.5 mV different from the Native case. F) PSPs predicted from conductances from example WT and Fmr1(-/y) cells. G) The predicted peak PSP for a population of WT and Fmr1(-/y) cells. * indicates significantly different means.

Next, we predicted PSPs from each cell we recorded from by running our model using the individual excitatory and inhibitory conductances measured from each of the WT and Fmr1(-/y) cells in our dataset. We found that overall, the predicted peak PSP is significantly reduced in Fmr1(-/y) cells compared to WT (**Fig. 4F-G**; n = 8 WT cells from 5 mice, 11 KO cells from 7 mice; Peak PSP WT = 16.25 ± 3.79 mV, Fmr1(-/y) = 9.69 ± 7.83 mV; p = 0.029). These results show that even though excitatory and inhibitory conductances scale up in Fmr1(-/y) cells compared to WT, the E-I balance limits the PSP amplitude to amplitudes at or below the WT amplitudes. The alteration in E-I balance in Fmr1(-/y) is even more extreme than what would be predicted by simply maintaining the WT PSP amplitudes. Overall, our results suggest that loss of Fmr1 may have differential effects on synaptic scaling in different brain areas, but that common compensatory mechanisms to stabilize post-synaptic potentials may be in place.

## DISCUSSION

Here we found that visual deficits observed in ASD may arise, at least in part, in the retina, at the earliest stages in visual processing. We find that sOn-a cells in Fmr1(-/y) retinas show changes in dendritic morphology and damped light responses that arise due to changes in E-I ratio.

Fmr protein (FMRP) regulates mRNA expression in dendrites, affecting key anatomical and physiological factors, especially at inhibitory synapses (Hagerman et al. 2017). sOn-α cell development in the mouse is characterized by the elaboration of their dendrites during the first two postnatal weeks (Lucas and Schmidt 2019). Here, we observed denser dendritic arbors and increased synaptic inhibition in the Fmr1(-/y) mouse, suggesting that both dendritic arbor development and synapse formation and/or pruning occur atypically. Denser dendrites and altered synaptic development have also been observed in a variety of other brain areas (Qin et al. 2011; He and Portera-Cailliau 2013), suggesting that FMRP may play similar roles in dendritic development in the retina as in other brain areas.

Altered synaptic signaling is a common theme across brain areas in Fragile X syndrome and also ASD more broadly (Coghlan et al. 2012; Contractor, Ethell, and Portera-Cailliau 2021). Changes in E-I ratio have been proposed as a common mechanism of dysfunction in ASD (Monday, Wang, and Feldman 2023), however more recent research in the cortex suggests that changes in E-I ratio are a compensatory mechanism to stabilize spiking (Antoine et al. 2019). Here, we found that synaptic changes in sOn-α cells are the opposite of what is observed in pyramidal cells in cortex in Fmr1(-/y) mice: both synaptic excitation and inhibition increase (rather than decrease in cortex) and E-I ratio goes down (rather than going up in cortex) (**Fig. 3**). However, these changes are still consistent with a model by Antoine et al. (Antoine et al. 2019) proposing that stabilized spiking requires non-linear scaling of excitation and inhibition, with the specific prediction that if excitation and inhibition increase, the E-I ratio should decrease, as was observed here (**Fig. 4**). Therefore, these results extend the validity of that model to cases where excitation and inhibition have increased. Understanding how synaptic and intrinsic properties of cells relate to their specific E-I balance and how this balance is established during development in sOn-α cells in the Fmr1 knockout will be important directions for future research.

Fmr1 is broadly expressed in the retina, not only in RGCs but also in the excitatory interneurons, the bipolar cells, and inhibitory interneurons, the amacrine cells (**Fig. 1, Fig. S1**). Within each of these cell classes, Fmr1 appears to be expressed at type-specific levels, suggesting that loss of Fmr1 could affect retinal cell types to a differing degree. Here, we found changes in the strength of synaptic input onto sOn-α cells and changes in dendritic density. Determining whether these differences occur due to loss of Fmr1 in sOn-α cells, in their presynaptic partners, or both will be an important next step to understanding how loss of Fmr1 affects retinal development.

How loss of Fmr1 will relate to visual symptoms in Fragile X syndrome is not yet obvious. Each RGC type may project to multiple brain areas (Kerschensteiner 2022), complicating inquiry into how changes in their signaling affect behavior. For example, sOn-α cells project to the superior colliculus, the lateral geniculate nucleus, and other targets. These cells provide information for behaviors including binocular-vision-guided hunting (Kim et al. 2020) and contrast sensitivity (Schmidt et al. 2014). How altered dim green vs. bright blue light sensitivity may influence the function of these circuits and their associated behaviors is not yet clear. Beyond the direct effects of lower light sensitivity on these circuits, altered activity of sOn-α cells during late visual development could influence development of downstream circuits in the brain through known activity-dependent mechanisms (Thompson et al. 2017).

Collectively, our results open up new avenues to understand the origin of sensory deficits in ASD. Further exploration of how different retinal cell types are affected by loss of Fmr1 and the downstream effects on brain and behavior will provide new insights into vision in Fragile X syndrome and ASD more broadly.

## Acknowledgements

Many thanks to Anis Contractor for providing Fmr1 knockout mice. Thanks to Greg Schwartz and Zach Jessen for providing the single-cell RNAseq dataset. The authors would like to acknowledge the staff of the Center for Comparative Medicine at Northwestern University for providing excellent animal care. Finally, thank you to the members of the Schmidt lab who provided training, support, and comments on the manuscript. This project received support from the Knights Templar Eye Foundation Career Starter Award (to AV); Northwestern SURG (to AW) and SIGP (to PG); NIH 5T32HL007909; and NIH R01EY030565 (to TS).

## Author contributions

(CRediT taxonomy, https://jats4r.org/credit-taxonomy)

Conceptualization: AV, TS; Data curation: AV, MS, PG, AW; Formal analysis: AV; Funding acquisition: AV, TS; Investigation: AV, MS, PG, AW; Methodology: AV; Project administration: TS; Resources: AV, MS, TS; Software: AV; Supervision: AV, TS; Validation: AV; Visualization: AV; Writing – original draft: AV; Writing – review and editing: all authors

## Declaration of interests

The authors have no conflicts of interest to declare.

## SUPPLEMENTAL 393 INFORMATION

**Figure S1.**
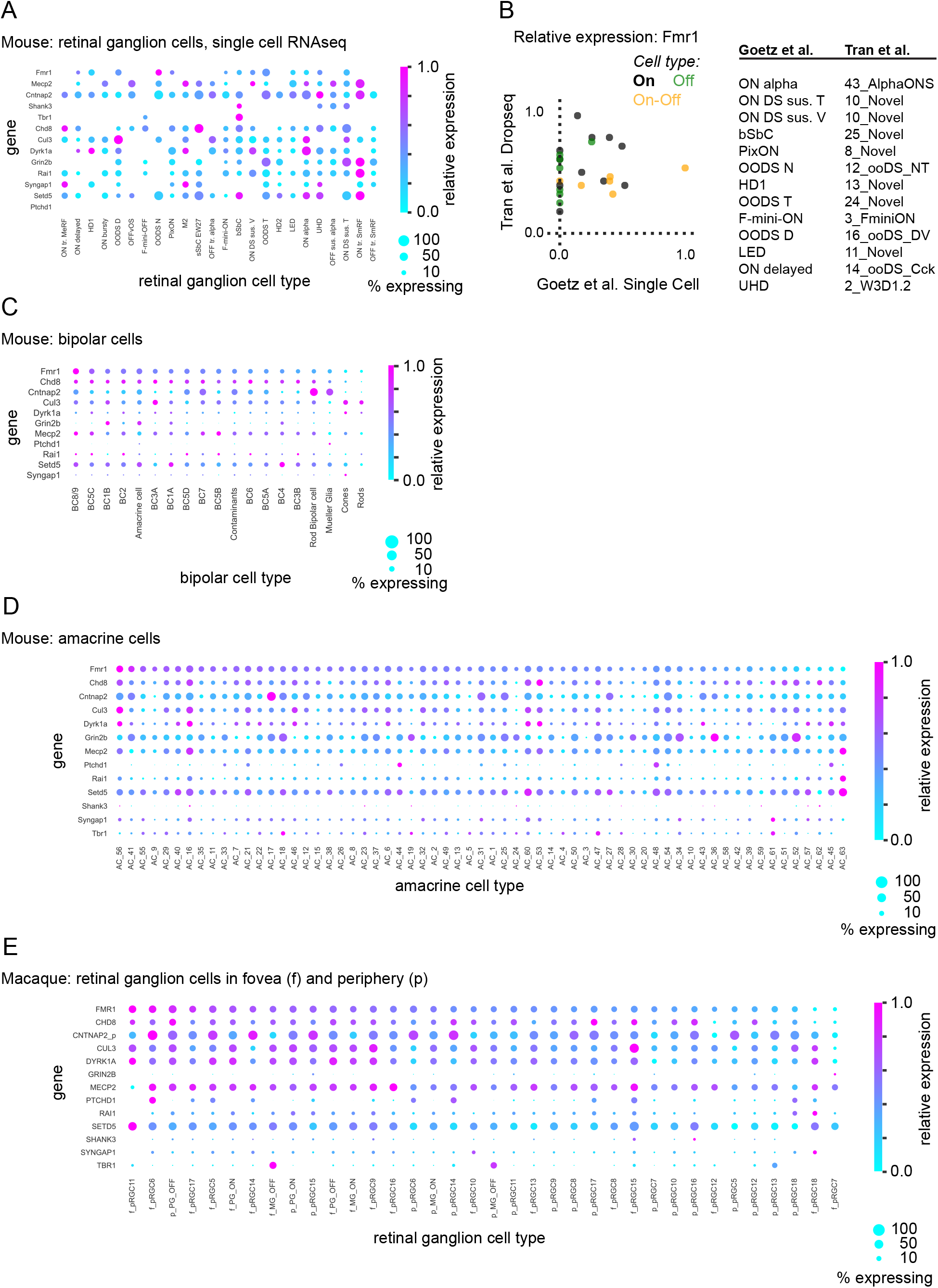
Autism-linked gene expression in the mouse and primate retina. Related to Figure 1. A) Type-specific expression of ASD-linked genes in mouse retinal ganglion cell types from a published single-cell RNAseq dataset (Goetz et al. 2022). Color: relative expression level within each gene. Dot size: percent of cells of each type expressing that gene. B) Left: Comparison of Fmr1 expression in two different mouse transcriptomes from **Fig. S1A** and Fig. 1A. Dots: RGC types’ relative Fmr1 expression in the Goetz et al. (2022) vs. Tran et al. (2019) datasets. Color indicates whether the cell type is in the On, Off or On-Off family. Right: list of cell types expressing Fmr1 in both datasets. Each row shows the cell type nomenclature for each dataset for matched cell types. C) Type-specific expression of ASD-linked genes in mouse bipolar cell types from a published bipolar cell transcriptome (Shekhar et al. 2016). D) Type-specific expression of ASD-linked genes in mouse amacrine cell types from a published amacrine cell transcriptome(Yan et al. 2020). E) Type-specific expression of ASD-linked genes in primate retinal ganglion cell types from a published retinal ganglion cell transcriptome (Peng et al. 2019).

**Figure S2.**
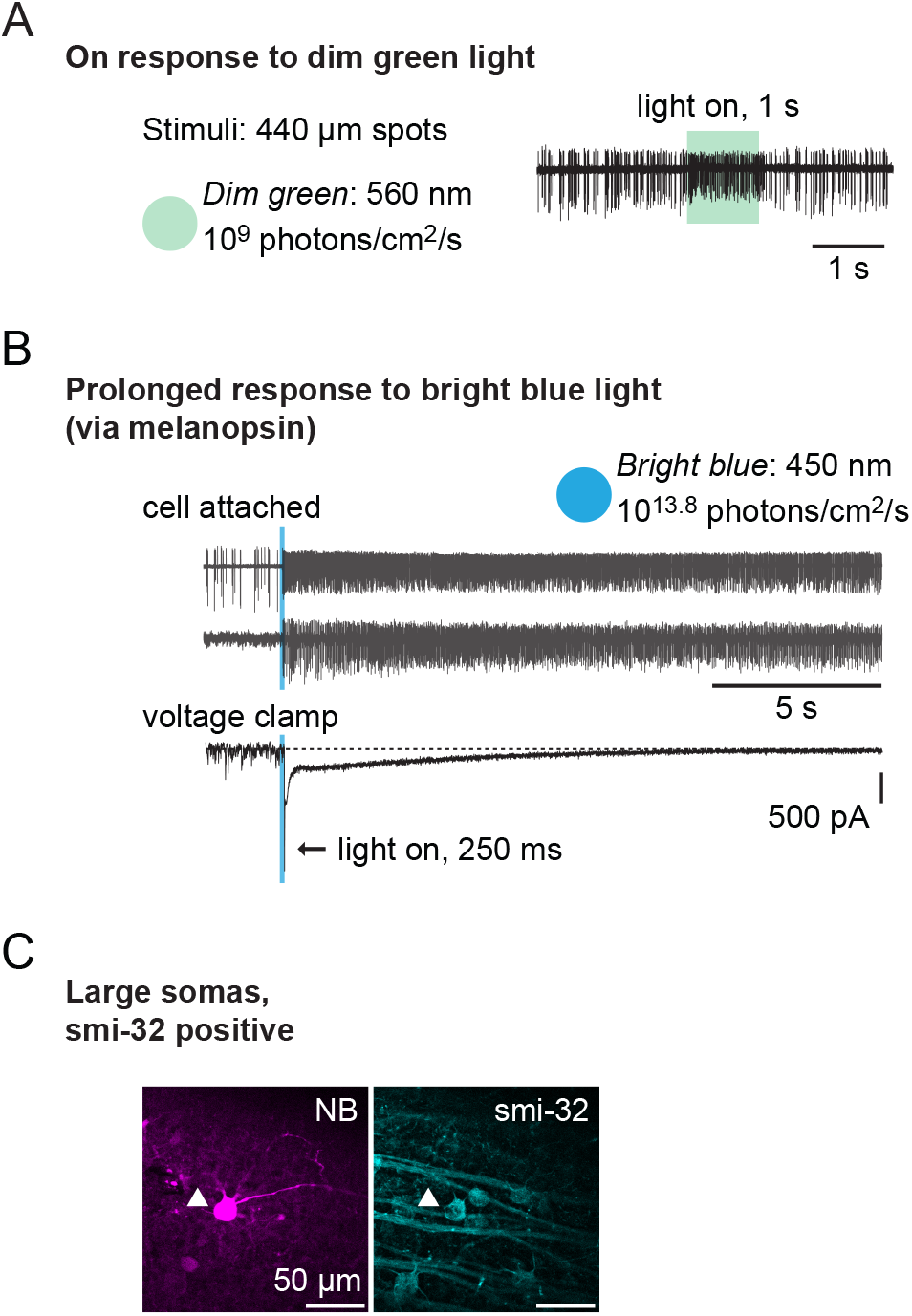
Identification of sOn-α cells. Related to Figures 1, 2. We identified putative sOn-α cells based on four features that, together, uniquely identify them. A) Cell attached electrophysiological recording from an sOn-α cell during presentation of a dim green light stimulus presented for 1 s. sOn-α cells exhibit characteristic high baseline firing rates and a sustained increase in firing when the light is turned on. Same data as in Fig. 2B. B) sOn-α cells are a type of intrinsically photosensitive melanopsin-positive ganglion cell. They exhibit prolonged firing in response to brief flashes of bright blue light, which optimally triggers the melanopsin expressed on their membranes. We checked for their melanopsin-dependent intrinsic photosensitivity by presenting a brief flash of bright blue light (blue rectangles) in either cell-attached (top two rows) to record the prolonged firing (note timescale) or voltage clamp (bottom row) to record the prolonged inward current. Recordings from three different cells are shown. C) sOn-α cells have large somas and are positive for smi-32. We filled recorded cells with neurobiotin and fixed and stained for smi-32. Here, a confocal image of an example sOn-α cell filled with neurobiotin (NB) and labeled with an antibody for smi-32.

**Figure S3.**
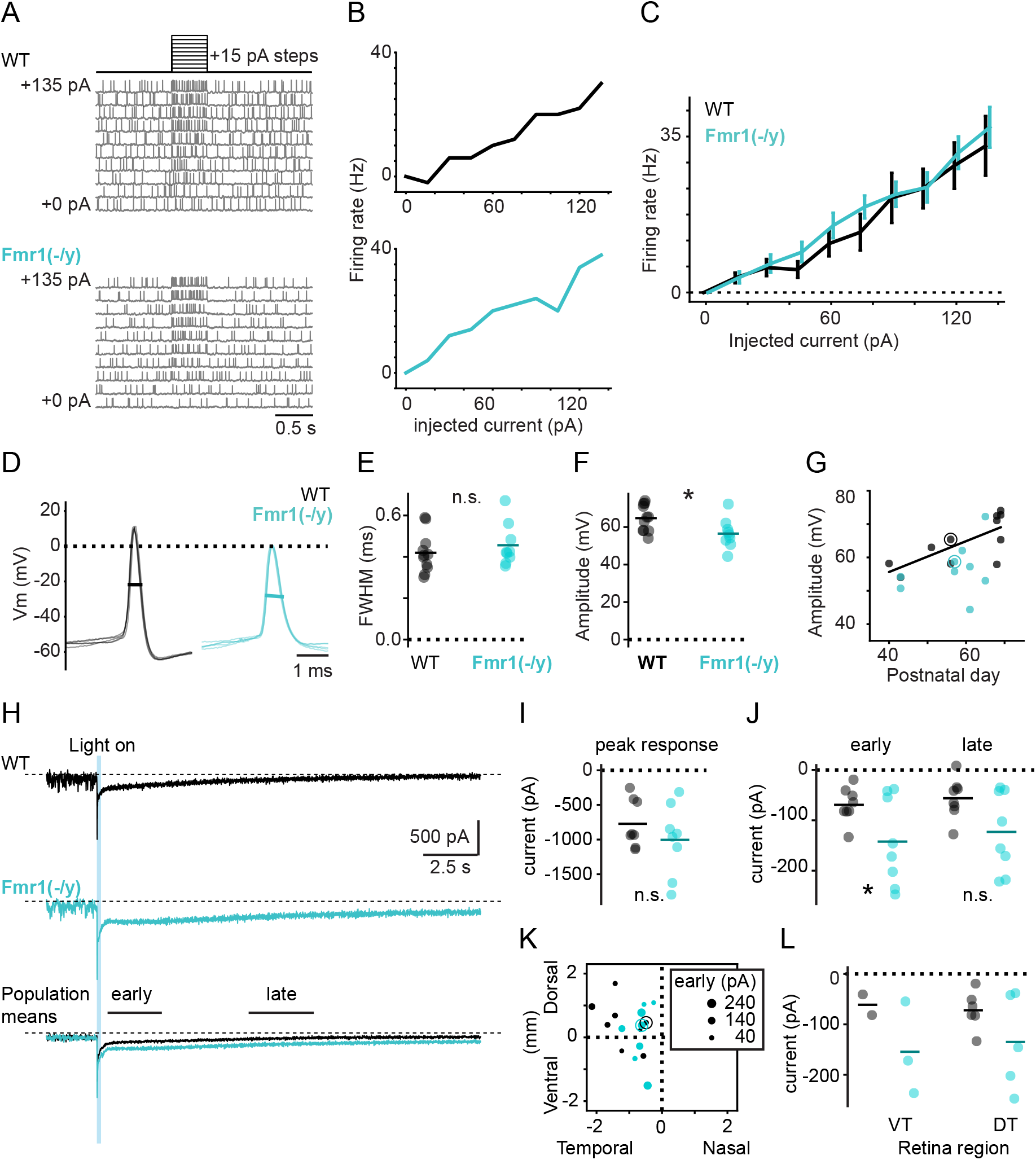
Subtle differences in intrinsic properties of sOn-α cells. Related to Figure 3. A) Current clamp recordings from two example sOn-α cells showing spiking in response to increasing current injections indicated by the protocol at the top. B) Firing rate as a function of the amount of current injected for the two example cells in **A**. C) Average firing rates as a function of current injected for sOn-α cells in WT and Fmr1(-/y) mice. Vertical lines: s.d. D) Example action potentials from two sOn-α cells. Multiple action potentials from each cell are overlaid. Vertical line indicates the full width half max (FWHM) measured in **E**. E) Average FWHM for sOn-α cells from WT and Fmr1(-/y) retinas. Population mean: lines. F) Spike amplitude from baseline for sOn-α cells from WT and Fmr1(-/y) mice. * indicates significant difference. G) Spike amplitude as a function of postnatal age of the animal. Black line: fit to the WT data. R-squared = 0.53. Circles: example cells from **D**. H) Voltage clamp recordings of bright blue stimulus-evoked currents in two example sOn-α cells (top, middle) and the population mean (bottom). Early and late epochs used for analysis are indicated. Stimulus period: blue rectangle. See also **Fig. S2**. I) Peak response during the stimulus period for sOn-α cells from WT and Fmr1(-/y) retinas. N.s. = not significant. Lines: population mean. J) Mean current change during early and late periods indicated in **H**. K) Mean current change during the early epoch (dot size) plotted on retina coordinates. L) Mean current change during the early epoch in the ventrotemporal (VT) and dorsotemporal (DT) regions of the retina.

**Figure S3 Results**: We tested whether excitability of sOn-α is altered in Fmr1(-/y) mice and found no difference between the current-spiking functions of WT vs. Fmr1(-/y) sOn-α cells (Sonoda et al. 2018) (**Fig. S3A-C**; n = 10 WT cells from 5 mice, 9 Fmr1(-/y) cells from 5 mice; two-way repeated measure ANOVA, p<0.01 for injected current, not significant for genotype or interaction). Second, we examined the properties of the action potentials themselves (**Fig. S3D-G**). While the spike width (full width half-max, FWHM, **Fig. S3E**) was not different between WT and Fmr1(-/y) retinas (WT: 0.419 ± 0.096 ms; Fmr1(-/y): 0.456 ± 0.101 ms; p = 0.422; n = 11 WT cells from 7 mice, 9 Fmr1(-/y) cells from 7 mice), the spike amplitude was slightly lower (**Fig. S3F**; WT: 64.8 ± 7.09 mV; Fmr1(-/y): 56.5 ± 7.77 mV; p = 0.025). In sOn-α cells, spike amplitude increases with developmental age (Lucas and Schmidt 2019). Here, we found that, while spike amplitude increases with age in WT mice, this relationship is less clear in the Fmr1(-/y) cells (**Fig. S3G**). Next, given that we found greater dendritic density in sOn-α cells in the Fmr1(-/y) mice (Fig. 1), we hypothesized that intrinsic properties that depend on membrane area could be atypical. sOn-α cells are intrinsically photosensitive retinal ganglion cells (type M4), so we wondered whether, the amount of melanopsin-mediated intrinsic photosensitivity in these cells could be increased given the cells’ longer dendritic lengths (Fig. 1). The melanopsin photocurrent is different from rod/cone evoked synaptic currents in that its duration is very prolonged, even when presented with a very short light pulse (**Fig. S1**) (Contreras et al. 2023). We observed that sOn-α cells in Fmr1(-/y) retinas had slightly larger, more prolonged blue light evoked currents (**Fig. S3H-J**, n = 8 WT cells from 6 mice, 8 KO cells from 6 mice; WT: -69 ± 34 pA, Fmr1(-/y): -142 ± 87 pA; p=0.045). However, this increased current in response to bright blue light cannot explain the lower firing rates in dimmer light conditions (Fig. 2).

## METHODS

### Animals

All procedures were approved by the Animal Care and Use Committee at Northwestern University. Male mice on a mixed B6/129 background were bred with female C57/Bl6J mice of either wildtype (WT) or Fmr1(-/-) (B6.129P2-Fmr1^tm1Cgr^/J, Jackson Labs # 003025)(Consortium 1994) genotypes to produce the WT and Fmr1(-/y) male animals used in this study. Tail biopsies were used to verify genotype. All mice were between P50-P75 except where otherwise noted.

### Retina dissection

All mice were dark adapted for at least one hour prior to being euthanized by CO_2_ asphyxiation followed by cervical dislocation. Under dim red light, the eyes were enucleated and retinas were dissected in carbogenated (95% O_2_, 5% CO_2_) Ames’ medium (Sigma-Aldrich or US Biological). Retinas were aligned using the subretinal vasculature(Wei, Elstrott, and Feller 2010) and cut into dorsotemporal and ventrotemporal sectors, which were mounted on a membrane filter (0.45 µm pore size, Millipore HABG01300) with a <1 mm^2^ hole cut in it. Retinas were maintained in carbongenated Ames media at room temperature for 30 minutes before transfer to the recording chamber.

### Electrophysiology

Electrophysiological recordings were performed using a previously described setup (Sonoda, Okabe, and Schmidt 2020). Retinas were perfused with carbogenated Ames media at 33-35° C and visualized using infrared illumination under DIC optics to minimize photobleaching of photoreceptors. sOn-α cells were identified by their large, square-ish cell somas (∼20 µm), sustained responses to dim green flashes, and prolonged responses to bright blue flashes (**Fig. 1**). Post-hoc staining and imaging confirmed alpha cell identity (see Immunohistochemistry) and stratification in the On layer relative to the ChAT bands. Boroscillate pipettes (Sutter Instruments, 3-5 MΩ) were used for all recordings and whole-cell electrophysiology recordings were performed using a Multiclamp 700B amplifier (Molecular devices). Data was acquired using a Digidata 1550B amplifier and data were collected using pClamp 10 acquisition software (Molecular Devices, RRID: SCR_011323).

For loose cell-attached recordings, pipettes were filled with Ames’ media and spikes were recorded in Multiclamp’s voltage clamp configuration, achieving a minimum seal resistance of at least 30 MΩ. For whole-cell current clamp recordings, the K^+^ internal contained: 125 K-gluconate, 2 CaCl_2_, 2 MgCl_2_, 10 EGTA, 10 HEPES, 2 ATP-Na_2_, 0.5 GTP-Na. For K^+^ internal, KOH was added to achieve pH of 7.22. For whole-cell voltage clamp recordings, the Cs^+^ internal contained (in mM): 110 CsMeSO_4_, 2.8 NaCl, 20 HEPES, 4 EGTA, 5 TEA-Cl, 4 ATP-Mg, 0.3 GTP-Na_3_, 10 Phosphocreatine-Na_2_, 5 QX-314-Br. For Cs^+^ internal, CsOH was added to achieve a pH of 7.2 and osmolarity was verified to be 290 mOsm/kg H_2_O. For both internal solutions, 0.3% neurobiotin and 10 µM Alexa-594 (ThermoFisher) were added to the internal solution for post-hoc visualization. For most experiments, one cell was recorded per retina piece to minimize contamination of light responses by repeat stimulation. For cell-attached recordings, if a cell responded to one dim green flash with a response other than a sustained On response (i.e. the cell was not a sustained On cell), the experimenter targeted a second cell in an area of the retina far from that site. Due to the time-consuming nature of blind-patching the alpha cells for electrophysiology, the electrophysiologist was not blinded to genotype during recording.

### Visual stimulation and recording protocols

A set of LED lights were used to stimulate the retina through the 60X water-immersion objective, achieving stimulus circle with a diameter of 440 µm. The shutter was controlled via pClamp and neutral density filters were used to control the light intensity. For the “dim green” stimulus, the wavelength was 560 nm, the irradiance was 10^9^ photons/cm^2^/s, and light was flashed for a duration of 1 s. For the “bright blue” stimulus, the wavelength was 450 nm, the irradiance was 10^13.8^ photons/cm^2^/s, and the light was flashed for a duration of 0.250 s. In most cases, both dim green and bright blue light responses were recorded from each cell. Dim green light was always presented first.

For voltage clamp recordings to collect data for conductance analysis, cells were held at 4 or 5 holding potentials spanning from the reversal potential for Cl to the reversal potential for cations and the light was flashed for 1 s at each holding potential. Data were acquired at 10 kHz and low-pass filtered at 2kHz. For bright blue light recordings, the cells were held at -73 mV. For current clamp recordings, a holding current was applied so that cells were resting at -60 mV, which typically required applying -50 to -100 pA of current.

### Immunohistochemistry

Retinas were fixed in 4% paraformaldehyde (Electron Microscopy Sciences) in PBS for 30 minutes at room temperature (RT), followed by three 30 minute washes in PBS at RT. Retinas were then blocked overnight at 4° C in PBS with 6% Normal Donkey Serum (NDS) and 0.3% Triton (blocking solution). Then retinas were incubated in blocking solution with primary antibodies for 3-4 nights at 4° C. Primary antibodies were 1:1000 streptavidin 546, 1:1000 mouse anti-smi-32, and 1:500 goat anti-ChAT. Next, retinas were washed three times for 30 minutes in PBS at RT, followed by overnight incubation at 4° C in blocking solution with secondary antibodies. Secondary antibodies were 1:1000 streptavidin 546, 1:1000 Alexa 647 donkey anti-mouse, and 1:500 Alexa 488 donkey anti-goat. Finally, retinas were washed 3x for 30 minutes in PBS and mounted on slides using Fluoromount (Sigma). Retinas were imaged on a confocal microscope (Leica DM5500 SPE, Leica Microsystems) under a 20X objective.

### Analysis

Analysis was performed using Excel (Microsoft) and Jupyter Notebook running Python (v 3.9.7). For plotting, the seaborn package was used (Waskom 2021). For statistical analysis, we used scipy (Virtanen et al. 2020) and pingouin (Vallat 2018) packages. Unless otherwise stated, reported p-values are the result of two-tailed student’s t tests.

#### Electrophysiology

Cells were discarded from analysis if they did not meet the criteria to be considered sOn-α cells, if they were not responsive to light, or if the access resistance changed during the recording or was higher than 50 MΩ. pClamp abf files were imported into Python using the pyABF package (Harden 2022). Custom scripts were used for spike detection, membrane property estimation, FWHM measurements, and conductance analysis (based on Vlasits et al. (2014)). Series resistance compensation was performed post-hoc for all voltage clamp recordings and voltages were corrected for the liquid junction potential (-10.5 mV).

E-I ratio was calculated as *G_e_/(G_i_+G_e_)* at each timepoint after conductances were denoised using a Savitzky-Golay filter. In some cases, the conductance values went below zero due to the high spontaneous baseline in retinal cells. Thus, we artificially shifted the baseline conductance by a factor of 4 to reduce the presence of negative values in the E-I ratio calculation.

#### Transcriptomic datasets

We collected a list of ASD-linked genes by searching PubMed for reviews on ASD-linked genes and from the SFARI list of mouse models of ASD (https://www.sfari.org/resource/mouse-models/). Most of the transcriptome datasets were accessed from singlecell.broadinstitute.org, except the single-cell RNAseq dataset, which was provided by that study’s authors (Goetz et al. 2022). Data for ASD-linked genes was imported into python using pandas (Reback and Team 2020) and plotted using seaborn.

#### Anatomical tracing and analysis

Analysis of anatomical data was performed by experimenters blinded to genotype. Confocal image stacks of neurobiotin-labeled sOn-α cells were imported into Fiji (Schindelin et al. 2012) for analysis. We used the SNT plugin to trace dendritic arbors, perform Sholl analysis, and measure the total dendritic length (Arshadi et al. 2021). The maximum radius and the dendritic diameter were manually measured in Fiji. To localize cells relative to the optic nerve, the coordinates of the optic nerve, cell, and cut edges of the retina were located and used to measure and align each retina piece using custom scripts in python.

#### Parallel conductance model

Light-evoked post-synaptic potentials were predicted with a parallel conductance model (Antoine et al. 2019), which we implemented in python using a jupyter notebook. The model predicted Vm using the parallel conductance equation:

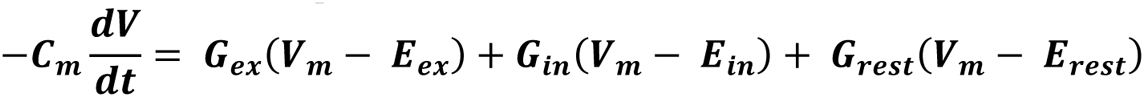

where C_m_ was the average membrane capacitance measured for either WT or Fmr1(-/y) cells (**Fig. 3**), E_ex_ was 0 mV, E_in_ was -72 mV, and G_rest_ was defined as the inverse of the average input resistance for either WT or Fmr1(-/y) cells (**Fig. 3**). E_rest_ was set to -55 mV to model the relatively depolarized state of sOn-α cells. For Fig. 4A-E, G_ex_ and G_in_ were the average conductances in WT cells. In Fig. 4F-G, each individual cell’s conductances were used.

Conductances were smoothed using a Savitzky-Golay filter and adjusted to remove negative values prior to use in the model. V_m_ was predicted using the forward Euler method.

## Notes

### Competing Interest Statement

The authors have declared no competing interest.

